# MMseqs2: sensitive protein sequence searching for the analysis of massive data sets

**DOI:** 10.1101/079681

**Authors:** Martin Steinegger, Johannes Söding

## Abstract

Sequencing costs have dropped much faster than Moore’s law in the past decade, and sensitive sequence searching has become the main bottleneck in the analysis of large metagenomic datasets. We therefore developed the open-source software MMseqs2 (mmseqs.org, which improves on current search tools over the full range of speed-sensitivity trade-off, achieving sensitivities better than PSI-BLAST at more than 400 times its speed.

Owed to the drop in sequencing costs by four orders of magnitude since 2007, many large-scale metagenomic projects each producing terabytes of sequences are being performed, with applications in medical, biotechnological, microbiological, and agricultural research^1;2;3;4^. A central step in the computational analysis is the annotation of open reading frames by searching for similar sequences in the databases from which to infer their functions. In metagenomics, computational costs now dominate sequencing costs^5;6;7^ and protein searches typically consume > 90% of computational resources^7^, even though the sensitive but slow BLAST^8^ has mostly been replaced by much faster search tools^9;10;11;12^. But the gains in speed are paid by lowered sensitivity. Because many species found in metagenomics and metatranscriptomics studies are not closely related to any organism with a well-annotated genome, the fraction of unannotatable sequences is often as high as 65% to 90%^13;2^, and the widening gap between sequencing and computational costs quickly aggravates this problem.

To address this challenge, we developed the parallelized, open-source software suite MMseqs2. Compared to its predecessor MMseqs^14^, it is much more sensitive, supports iterative profile-to-sequence and sequence-to-profile searches and offers much enhanced functionality (**Supplementary Table S I**).

MMseqs2 searching is composed of three stages (Fig. 1a): a short word (“*k*-mer”) match stage, vectorized ungapped alignment, and gapped (Smith-Waterman) alignment. The first stage is crucial for the improved performance. For a given query sequence, it finds all target sequences that have two consecutive similar-*k*-mer matches on the same diagonal (Fig. 1b). Consecutive *k*-mer matches often lie on the same diagonal for homologous sequences (if no alignment gap occurs between them) but are unlikely to do so by chance. Whereas most fast tools detect only *exact k*-mer matches^9;10;11;12^, MMseqs2, like MMseqs and BLAST, finds *k*-mer matches between *similar k*-mers. This similar-*k*-mer matching allows MMseqs2 to use a large word size *k*=7 without loosing sensitivity, by generating a large number of similar *k*-mers, ∼600 to 60 000 per query *k*-mer depending on the similarity setting (Fig. 1b, orange frame). Importantly, its innermost loop 4 needs only a few CPU clock cycles per *k*-mer match using a trick to eliminate random memory access (last line in magenta frame, **Supplementary Fig. S1**).

**Figure 1:**
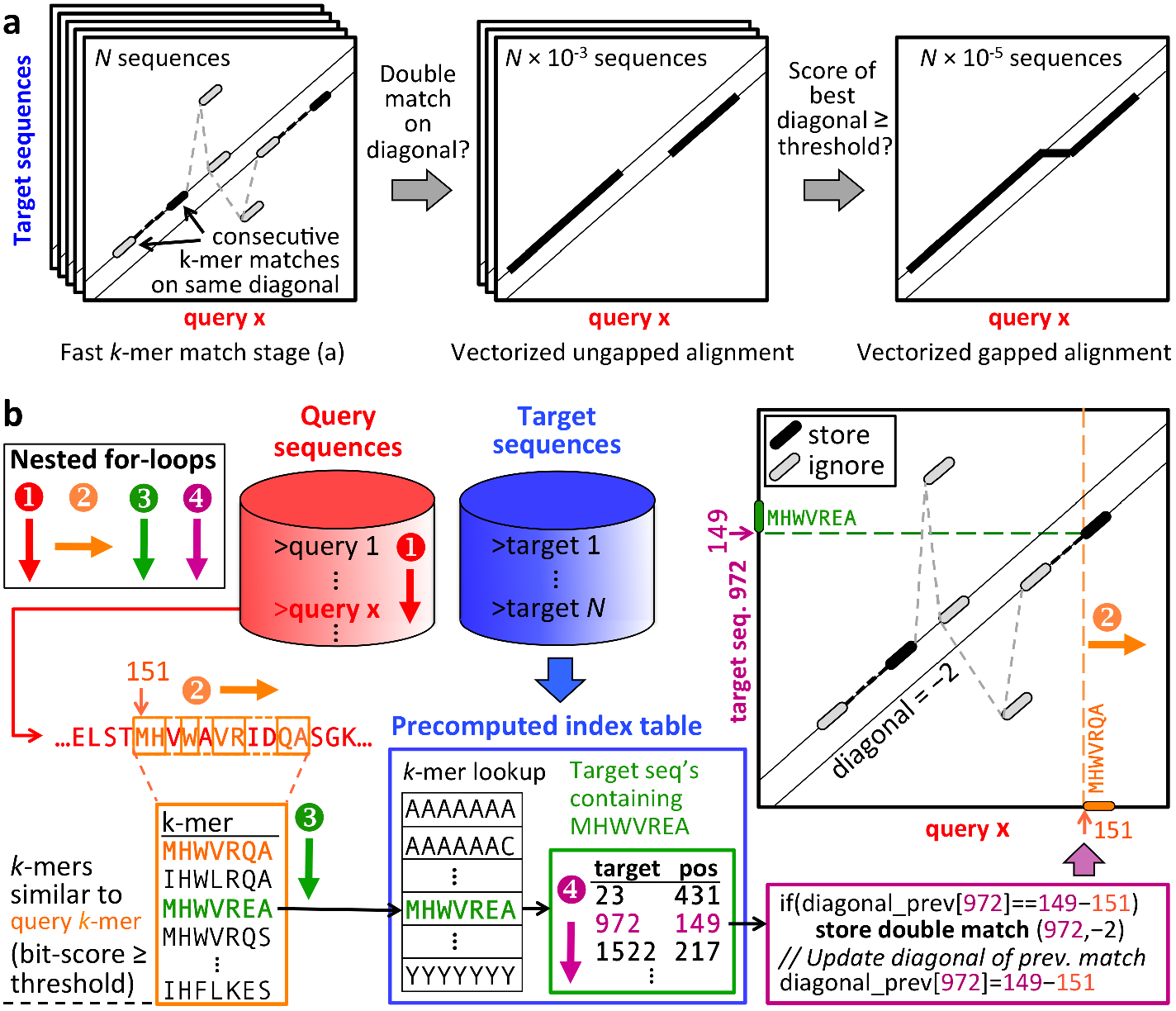
MMseqs2 searching in a nutshell. **(a)** Three increasingly sensitive search stages find similar sequences in the target database. (b) The short word (”*k*-mer”) match stage detects consecutive inexact *k*-mer matches occurring on the same diagonal. The diagonal of a *k*-mer match is the difference between the positions of the two similar *k*-mers in the query and in the target sequence. The pre-computed index table for the target database (blue frame) contains for each possible *k*-mer the list of the target sequences and positions where the *k*-mer occurs (green frame). Query sequences/profiles are processed one by one (loop 1). For each overlapping, spaced query *k*-mer (loop 2), a list of all similar *k*-mers is generated (orange frame). The similarity threshold determines the list length and sets the trade-off between speed and sensitivity. For each similar *k*-mer (loop 3) we look up the list of sequences and positions where it occurs (green frame). In loop 4 we detect consecutive double matches on the same diagonals (magenta and black frames).

The critical insight was to follow our credo “information is power” by combing the double-match criterion with making *k*-mers as long as possible, which required finding similar and not just exact *k*-mers. This effectively bases our decision on up to 2×7=14 residues instead of just 2 × 3 in BLAST or 12 letters on a size-11 alphabet in DIAMOND.

MMseqs2 is parallelized on three levels: time-critical parts are manually vectorized, queries can be distributed to multiple cores, and the target database can be split into chunks distributed to multiple servers. Because MMseqs2 needs no random memory access in its innermost loop, its runtime scales almost inversely with the number of cores used (**Supplementary Fig. S2**).

MMseqs2 requires 13.4 GB plus 7B per amino acid to store the database in memory, or 80 GB for 30.3 M sequences of length 342. Large databases can be searched with limited main memory by splitting the database among servers, at very moderate loss of speed (**Supplementary Fig. S3**).

We developed a benchmark with full-length sequences containing disordered, low-complexity and repeat regions, because these regions are known to cause false-positive matches, particularly in iterative profile searches. We annotated UniProt sequences with structural domain annotations from SCOP^15^, 6370 of which were designated as query sequences and 3.4 M as database sequences. We also added 27 M reversed UniProt sequences, thereby preserving low complexity and repeat structure^16^. The unmatched parts of query sequences were scrambled in a way that conserved the local amino acid composition. A benchmark using only unscrambled sequences gives similar results (**Supplementary Figs. S4, S5, S6, S7**). We defined true positive matches to have annotated SCOP domains from the same SCOP family, false positives match a reversed sequence or a sequence with a SCOP domain from a different fold. Other cases are ignored.

Figure 2a shows the cumulative distribution of search sensitivities. Sensitivity for a single search is measured by the area under the curve (AUC) before the first false positive match, i.e., the fraction of true positive matches found with better *E*-value than the first false positive match. MMseqs2-sensitive reaches BLAST’s sensitivity while being 36 times faster. Interestingly, MMseqs2 is as sensitive as the exact Smith-Waterman aligner SWIPE^17^, compensating some unavoidable loss of sensitivity due to its heuristic prefilters by effectively suppressing false positive matches between locally biased segments (Fig. 2d, **Supplementary Fig. S4**). This is achieved by correcting the scores of regions with biased amino acid composition or repeats, masking such regions in the *k*-mer index using TANTAN^18^, and reducing homologous overextension of alignments^19^ with a small negative score offset (Fig. 2d, **Supplementary Fig. S7**). All tools except MMseqs2 and LAST reported far too optimistic *E*-values (**Supplementary Fig. S8**). For example in the 6370 searches DIAMOND reported 69211 false positive matches with *E*-values below 10^−3^ (versus 0.637 expected) in 5% of the searches (versus 0.1% expected), while MMseqs2 produced 54 false positive matches with *E* < 10^−3^ in only 0.1% of the searches (Supplemental Table S1). In automatic functional annotation pipelines, such unreliable *E*-values will lead to an increased fraction of false annotations.

**Figure 2:**
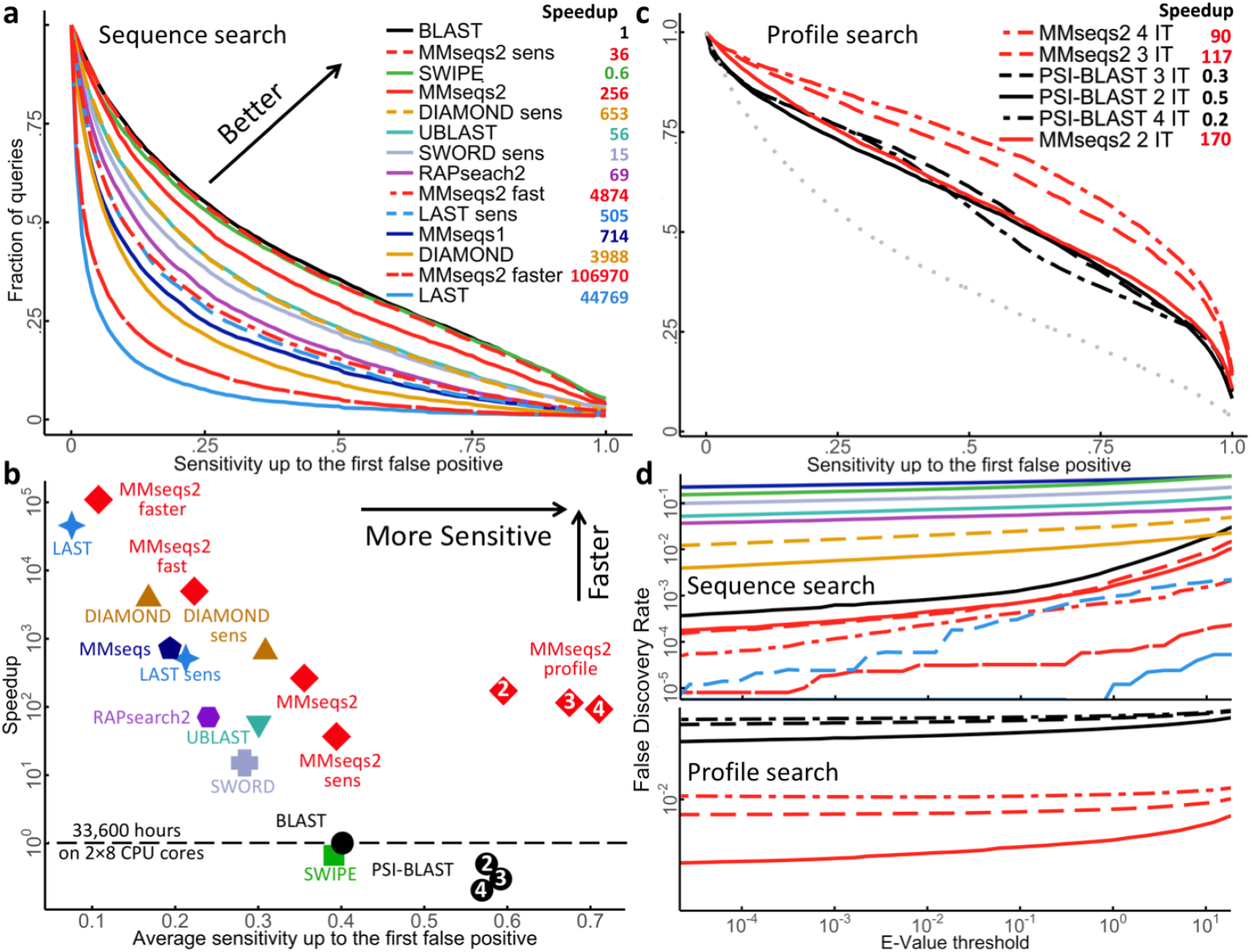
MMseqs2 pushes the boundaries of sensitivity-speed trade-off. **a** Cumulative distribution of Area under the curve (AUC) sensitivity for all 6370 searches with UniProt sequences through the database of 30.4 M full-length sequences. Higher curves signify higher sensitivity. Legend: speed-up factors relative to BLAST, measured on a 2 × 8 core 128 GB RAM server using a 100 times duplicated query set (637 000 sequences). Times to index the database have not been included. MMseqs2 indexing takes 11 minutes for 30.3M sequences of avg. length 342. **b** Average AUC sensitivity versus speed-up factor relative to BLAST. White numbers in plot symbols: number of search iterations. **c** Same analysis as in a, for iterative profile searches. **d** False discovery rates for sequence and profile searches. Colors: as in a (top) and c (bottom).

In a comparison of AUC sensitivity and speed (Fig. 2b), MMseqs2 with four sensitivity settings (red) shows the best combination of speed and sensitivity over the entire range of sensitivities. Similar results were obtained with a benchmark using unscrambled or single-domain query sequences (**Supplementary Figs. S4, S5, S6, S7, S9, S10**).

Searches with sequence profiles are generally much more sensitive than simple sequence searches, because profiles contain detailed, family-specific preferences for each amino acid at each position. We compared i MMseqs2 to PSI-BLAST (Fig. 2b) using two to four iterations of profile searches through the target database. As expected, MMseqs2 profile searches are much faster and more sensitive than BLAST sequence searches. But MMseqs2 is also considerably more sensitive than PSI-BLAST, despite being 433 times faster at 3 iterations. This is partly owed to its effective suppression of high-scoring false positives and more accurate *E*-values (Fig. 2d, **Supplementary Fig. S7**).

The MMseqs2 suite offers workflows for various standard use cases of sequence and profile searching and clustering of huge sequence datasets and includes many utility scripts. We illustrate its power with three example applications.

In the first example, we tested MMseqs2 for annotating proteins in the Ocean Microbiome Reference Gene Catalog (OM-RGC)^1^. The speed and quality bottleneck is the search through the eggNOGv3 database^20^. The BLAST search with *E*-value cutoff 0.01 produced matches for 67% of the 40.2 M OM-RGC genes^1^. We replaced BLAST with three MMseqs2 searches of increasing sensitivity (**Supplementary Fig. S11**). The first MMseqs2 search in fast mode detected matches for 59.3% of genes at *E* ≤ 0.1. (*E* ≤ 0.1 corresponds to the same false discovery rate as *E* ≤ 0.01 in BLAST, Fig. 2d). The sequences without matches were searched with default sensitivity, and 17.5% had a significant match. The last search in sensitive search mode found matches for 8.3% of the remaining sequences. In total we obtained at least one match for 69% sequences in OM-RGC, 3% more than BLAST, in 1% of the time (1 520 vs. 162 952 CPU hours; Shini Sunagawa, personal communication).

In the second example, we sought to annotate the remaining 12.3 M unannotated sequences using profile searches. We merged the UniProt database with the OM-RGC sequences and clustered this set with MMseqs2 at 50% sequence identity cut-off. We built a sequence profile for each remaining OM-RGC sequence by searching through this clustered database and accepting all matches with *E* ≤ 0.001. With the resulting sequence profiles we searched through eggNOG, and 3.5 M (28.3%) profiles obtained at least one match with *E* < 0.1. This increased the fraction of OM-RGC sequences with significant eggNOG matches to 78% with an additional CPU time of 900 hours. In summary, MMseqs2 matched 78%) sequences to eggNOG in only 1.5% of the CPU time that BLAST needed to find matches for 67% of the OM-RGC sequences^1^.

In the third example, we annotated a non-redundant set of 1.1 billion hypothetical j proteins sequences with Pfam^21^ domains. We predicted these sequences of average length 134 in ∼2200 metagenome/metatranscriptome datasets^22^. Each sequence was searched through 16 479) Pfam31.0 sequence profiles held in 16 GB of memory of a single 2 × 14-core server using sensitivity setting `-s 5`. **Supplementary Fig. S12** explains the adaptations to the *k*-mer prefilter and search workflow. The entire search took 8.3 hours, or 0.76 ms per query sequence per core and resulted in 370 M domain annotations with *E*-values below 0.001. A search of 1100 randomly sampled sequences from the same set with HMMER3^23^ through Pfam took 10.6 s per seqeunce per core, almost 14 000 times longer, and resulted in 514 annotations with *E* < 0.001, in comparison to 415 annotations found by MMseq2. A sensitivity setting of j `-s 7` brings the number of MMseqs2 annotations to 474 at 4000 times the speed of HMMER3.

In summary, MMseqs2 closes the cost and performance gap between sequencing and computational analysis of protein sequences. Its sizeable gains in speed and sensitivity should open up new possibilities for analysing large data sets or even the entire genomic and metagenomic protein sequence space at once.

## ACKNOWLEDGEMENTS

We are grateful to Cedric Notredame and Chaok Seok for hosting M.S. at the CRG in Barcelona and at Seoul National University for 12 and 18 months, respectively, and to Burkhard Rost at TU Munich for accepting the formal supervision of his PhD thesis. We thank Milot Mirdita, Lars van den Driesch and Clovis Galiez for contributing utilities and workflows, and Shini Sunagawa, Martin Frith, Thomas Rattei and our lab for feedback on the manuscript. This work was supported by the European Research Council’s Horizon 2020 Framework Programme for Research and Innovation (‘’Virus-X”, project no. 685778) and by the German Federal Ministry for Education and Research (BMBF) (grants e:AtheroSysMed 01ZX1313D, “SysCore” 0316176A).

## AUTHOR CONTRIBUTIONS

M.S. developed the software and performed the data analysis. M.S. and J.S. conceived of and designed the algorithms and benchmarks and wrote the manuscript.

